# Over-the-counter fiber supplementation associates with metabolic and microbial shifts in rectal mucosa

**DOI:** 10.64898/2026.09.08.750137

**Authors:** David A. Alvarado, Lydia Okyere, Amber O’Connor, Maura Walsh, Siting Chen, Vassiliki Liana Tsikitis, Christopher Gaulke

## Abstract

Fiber supplements are the first line of treatment for all patients with benign anorectal disease. In addition, fiber has been found to support gut microbiota and reduce inflammation; however, their integrated effect on the colonic mucosal environment is incompletely defined. We evaluated how mixed-fiber formulations may collectively shape the rectal mucosal environment through coordinated effects on microbiota composition, short-chain fatty acid (SCFA) handling, and host transcription. We evaluated multi-compartment responses to a 28-day mixed-fiber intervention in 37 participants with benign anorectal disease (∼15 g/day; 7.5g psyllium husk, 8g wheat dextrin) and assessed the microbiota (16S rRNA), the metabolome (GC-MS and LC-MS), and transcriptome (subset n=10) before (PRE) and after (POST) supplementation. Changes in KEGG pathways were tested with gene set enrichment analysis (GSEA). Associations were tested by Spearman correlation, and microbiota-based prediction of SCFA responses by random-forest regression. Microbiota diversity was stable after the fiber intervention, however, 10 mucosal ASVs were differentially abundant including *Parabacteroides* and *Lachnospiraceae* NK4A136 group while SCFAs, propionate and isobutyrate, both decreased in rectal tissue and butyrate decreased in serum. To understand these shifts in the metabolome, we proceeded with the transcriptome where we identified 130 differentially expressed genes and enrichment of metabolic and SCFA-related pathways. Random forest regression captured a modest signal for serum butyrate. Taken together, short-term mixed-fiber supplementation produced selective rectal mucosal taxonomic shifts and enriched mucosal metabolic and SCFA-metabolism program despite reduced SCFA pools, suggesting that even brief over-the-counter fiber supplementation may reshape the mucosal metabolic environment.

**Importance:** Dietary fiber plays a critical role in shaping gut microbial composition and function. While impacts of fibers on the fecal microbiome have been extensively studied, less is known about their effects on mucosal populations. Mucosal microbial communities are distinct from fecal microbiota and play an important role in barrier function, inflammation, and gut health. To better understand how fiber may impact these communities we quantified changes mucosal microbiome composition and metabolism before and after a 28-day over-the-counter fiber supplementation. Unlike previous work in feces, fiber supplementation did not result in broad mucosal community restructuring but rather modest specific taxonomic shifts and alterations in microbial metabolism with concomitant reduced inflammatory and increased SCFA-utilization gene expression in tissue. These niche specific differences in responses to fiber underscore the need for further study of the role of the mucosal microbiome in human health.

## Introduction

Benign anorectal diseases encompass a diverse spectrum of prevalent structural and functional conditions such as hemorrhoids, anal fissures, anorectal abscesses, fistulas, defecation dyssynergia, and fecal incontinence, that manifest with distressing symptoms including bleeding, pain, infection/drainage, and impaired continence ^1–4^. The population burden is substantial, driven largely by hemorrhoids and anal fissures. Hemorrhoids affect 10 million U.S. adults annually and are observed in up to 38% of screening colonoscopies ^1,2^ and anal fissures carry an estimated 11% lifetime prevalence ^1–3,5,6^. Collectively, benign anorectal disease accounts for up to 20% of all outpatient surgical referrals ^7^, and severity ranges from self-limiting discomfort to profound functional impairment and, in rare cases, life-threatening perineal sepsis or gangrene ^1–3^. These conditions carry major quality-of-life and socioeconomic consequences, including psychological distress and loss of independence among older adults ^4,7^. Delayed or unsupervised care often driven by stigma and access barriers further increases morbidity through complications such as severe bleeding, stenosis, and, in extreme cases, permanent colostomy ^7^.

Dietary fiber is widely recommended as a foundational, first-line treatment for multiple benign anorectal conditions, as it significantly improves overall patient quality of life ^6^ by improving stool consistency, reducing straining, and relieving symptoms. Specifically, adequate fiber intake has been shown to consistently reduce bleeding and overall symptoms in hemorrhoids, lower the recurrence rate of anal fissures, and aid in the daily management of both rectal pain and bleeding ^3,6,8^. As defined by the FDA, dietary fibers are non-digestible soluble and insoluble carbohydrates with ≥3 monomeric units (e.g., resistant maltodextrin/dextrin), and lignin that are intrinsic in plants, or isolated and synthetic non-digestible carbohydrates (e.g., psyllium husk) determined to have a physiological effect that are beneficial to human health ^9,10^. Contemporary guidance recommends dietary and behavioral modification as initial therapy for symptomatic hemorrhoids, and for anal fissures a total fiber intake of about 25–35 g/day using diet and over-the-counter supplementation ^6,8^. A systematic review and meta-analysis demonstrated that fiber supplementation reduced the risk of persistent hemorrhoidal symptoms by 47% and bleeding by 50%, supporting its role as an evidence-based conservative treatment ^8,11,12^.

A standard fiber regimen prescribed for benign anorectal disease targets 20–35 g/day paired with 1.5–2 L of water, with the goal of producing a daily soft stool without straining ^6^. To achieve this, physicians frequently prescribe supplements such as psyllium at 7–20 g ^1–3,12^. Both psyllium husk and wheat dextrin are FDA-recognized dietary fibers available as common over-the-counter supplements ^10^. Wheat dextrin is fully soluble and fermentable; made up of branched α-1,2 and α-1,3 linkages generated during dextrinization which confer resistance to small intestinal digestion, and its fiber benefits are reported for glucose and cholesterol regulation ^10^. Reported by the European Food Safety Authority (EFSA), wheat dextrin health claims are related to the maintenance of normal blood pressure, normal (fasting) blood concentrations, reduction of post-prandial glycemic response, increase in magnesium and/or calcium retention, and normal bowel function ^20^. Psyllium husk is also primarily soluble (60–80%) and slowly fermentable; its principal component arabinoxylan is composed of β-1,4, α-1,2, α-1,3, and O-5 ester linkages, resists small intestinal digestion and has been reported to improve gastrointestinal health through stool bulking in addition to glucose and cholesterol regulation ^10,21,22^*. In vitro* fermentation demonstrated that wheat dextrin can be utilized by the microbes with the CAZymes to break down the branched α-linkages ^23^, while psyllium, due to the formation of the gel and its insoluble properties, requires specialized primary degraders to metabolize the polymer and release fragments or metabolites that can support cross-feeding by secondary consumers ^24–26^. Dietary fibers that escape host digestion in the small intestine are degraded by the colonic microbiota into short-chain fatty acids (SCFAs), principally acetate, propionate, and butyrate, as major functional outputs of fiber–microbiome interactions ^25,27,28^. Among these, butyrate is particularly relevant to colorectal health because it supports colonocyte metabolism, mucosal integrity, and local immune regulation, and has been linked to protection from neoplastic change ^2,29,30^.

Fiber-rich diets promote the growth of beneficial fiber-degrading microbes that help maintain barrier function and stimulate mucus secretion ^13–15^. Conversely, fiber deprivation may drive the gut microbiota to degrade host-secreted mucus glycoproteins as an alternative nutrient source, thinning the protective mucosal barrier and increasing pathogen susceptibility ^14,15^. A high-fiber-dose, inulin-type fructan, has been shown to impact the mucosal microbiome differently than the fecal microbiome, targeting specific opportunistic pathogens ^13^, and reducing tissue inflammation ^16^. Furthermore, prospective clinical evidence underscores the diagnostic superiority of the mucosal environment, demonstrating that the brush-border associated microbiota is a far more sensitive indicator of health than intraluminal populations, accurately predicting adenoma status outperforming paired fecal and oral samples ^17^. Clinically, nearly all human evidence linking fiber to the gut microbiota derives from fecal sampling, which underplays the mucosa-associated community in direct contact with the epithelium most relevant to barrier integrity, bleeding, and inflammation in benign anorectal disease ^13,16–19^. Although the rectal mucosal community is compositionally variable from the fecal community and a more sensitive marker of mucosal healthy, whether routine over-the-counter fiber supplementation remodels this mucosa-associated community and its local metabolome remains poorly characterized.

Building on these observations, we leveraged a one-month standard-of-care fiber intervention for benign anorectal disease to determine whether routine over-the-counter fiber supplementation could measurably shift the rectal mucosal microbiota and metabolome. We applied a multi-omic methodology with a paired, within-subject design (PRE-vs POST-supplementation) using a cohort of patients undergoing standard-of-care treatment with a mixed soluble and insoluble fiber preparation, chosen because it is easy to obtain, feasible to administer, and already routinely used in clinical practice. This allowed us to study a pragmatic, real-world intervention while examining whether mixed-fiber supplementation influences the rectal mucosal microbiota and metabolome beyond symptom control.

## Methods

### Study design

This study evaluated a 28-day fiber supplementation intervention consisting of 15.5 g of total daily fiber, administered as 2.5 packets of Benefiber On the Go Prebiotic Fiber Supplement (7.5 g fiber) and 2 packets of Metamucil Fiber Thins (8 g fiber). Study visits were conducted at Oregon Health & Science University (OHSU) and included collection of demographic information and questionnaires assessing physical activity, habitual diet, and medical history. Patients were contacted weekly via telephone to assess fiber compliance and adherence to study protocol. A total of 110 participants were enrolled at OHSU and provided written informed consent before undergoing any study procedures. Participants were instructed to maintain their usual diet and lifestyle during the intervention period while adhering to the prescribed fiber supplementation regimen. Self-reported antibiotic use within the 90 days prior to intervention (n=7) was not an exclusion criterion, as the study was designed to reflect a clinically heterogeneous real-world population. This study was reviewed and approved by the Oregon Health & Science University (OHSU) Institutional Review Board (IRB Protocol #24054) and conducted following the principles of the Declaration of Helsinki. From the total enrolled participants (110), 38 participants were excluded prior to intervention (e.g., did not meet inclusion criteria at time of appointment, no show to study visit or discontinued due to participant discretion) or screening failures. Of the 72 participants that provided PRE-samples, 24 patients were lost to follow-up (e.g., did not return for study visit 2, fiber-related discomfort or discontinued due to participant discretion). Of the 48 paired samples collected, yielded a final analytic cohort of 37 per-protocol, defined as a POST biopsy collected within 7 days of the final fiber dose date (**Supplemental Figure 1**).

### Sample processing

Rectal mucosal biopsies were obtained with Boston scientific cold forceps from participants during proctoscopy at their first visit (PRE) and at the end of the 28-day intervention (POST), along with paired blood serum samples. While the serum samples were collected in EDTA tubes and centrifuged to separate plasma, biopsies were flash-frozen in liquid nitrogen and both were stored at -80C until processing. Of the participants that could provide samples, a total of 72 PRE-samples and 48 paired POST-samples were collected. The RNA and microbial DNA from rectal biopsies were extracted using the AllPrep DNA/RNA kit (QIAGEN) as previously described ^17^. Each samples’ DNA was quantified using a Qubit3.0 Fluorometer (ThermoFischer Scientific) before sending to the Roy J. Carver Biotechnology Center. 16S amplicons were generated with V4 primers (515F: 5’-GTGYCAGCMGCCGCGGTAA, 806R: 5’-GGACTACNVGGGTWTCTAAT) ^31^ and sequenced on an Illumina MiSeq i100 25M flowcell generating 2×300nt paired-end reads.

### Microbial Community Diversity and Differential Abundance Analyses

Raw sequence data were processed using the ‘DADA2’ (v1.30) pipeline ^32^ in R (v4.4.1). Reads were quality-filtered (*truncLen*=c(205, 247), *maxN*=0, *maxEE*=c(2,2), *truncQ*=2, and *rm.phix*=T), denoised, merged and filtered of chimeric sequences (*removeBimeraDenovo*, method=“consensus”). Taxonomic classification was performed against the ‘SILVA’ (v138.1) reference database using *assignTaxonomy* ^33^. Only ASVs classified as Bacteria were retained for downstream analyses, removing 34 non-bacterial ASVs to yield a final dataset of 9,110 bacterial ASVs. Sequencing depth of the ASV table was normalized by rarefaction to 6,500 reads per sample using ‘vegan’ (v2.6-8; *rrarefy*, *seed*=8), based on the minimum library size (6,603 reads). After rarefaction, ASVs with zero counts across all samples and singletons were filtered out.

Richness (number of unique ASVs per sample) and Shannon index (accounting for both richness and evenness) was calculated with ‘vegan’ (*specnumber* and *diversity*, respectively). Beta diversity was quantified using Jaccard dissimilarity (*vegdist*, *method*=“jaccard”). Non-metric multidimensional scaling (NMDS) ordination was performed with ‘vegan’ (*metaMDS; distance*=“jaccard”, *k*=3) resulting in stress=0.143, as k=2 produced higher stress (0.204). Prior to differential abundance analysis, commonly applied prevalence and minimum-abundance heuristics ^34^ to reduce sparsity and improve stability of downstream inference. ASVs were required to be detected in ≥10% of samples and reach a minimum abundance of ≥0.1% reads.

### Targeted metabolomics

Tissue samples were homogenized, extracted with 70% methanol and subjected to SCFA analysis first. Serum samples were spiked with pure methanol for deproteinization, centrifuged 10 min at 4C (16,000 rpm), spiked with internal standard (13C Acetate) and analyzed for SCFA. Technical batch effects in serum and tissue metabolites concentrations were evaluated and corrected prior to analysis. Tissue and serum metabolites were analyzed separately. Tissue VFAs included zeros (values at/under detection limit) and were handled by adding a pseudocount equal to half the minimum non-zero value prior to log transformation; serum SCFAs had no zero values and were log-transformed directly ^35,36^. Additive batch effects were removed using ‘limma’ v3.66.0 (*removeBatchEffect*) ^37^, preserving the timepoint contrasts, and verified by batch association in corrected values (all *P*>0.80).

### Tissue bulk RNA-seq

Rectal mucosal biopsy RNA-seq was performed on a subset of the cohort (N=10; paired) based on availability and quality of biopsies. Library preparation and sequencing were conducted at the Roy J. Carver Biotechnology Center DNA Services using the Agilent SureSelect XT HS2 kit. Libraries were pooled, quantified by qPCR, and sequenced on an Illumina NovaSeq X Plus to generate paired-end reads (2×151). Reads were adapter and quality trimmed with ‘fastp’ (v1.0.1) prior to alignment. Libraries were strand-specific (Read 1 aligned to antisense strand; Read 2 aligned to sense strand) and trimmed reads were aligned to the human reference genome (GRCh38 primary assembly, ‘Ensembl release 115’) using ‘STAR’ (v2.7.11b) ^38^. Gene-level read summarization was performed using ‘featureCounts’ ^39^ to enable explicit counting parameters including paired-end mode, reverse-stranded counting (-s 2), summarized (-t exon), and aggregated (-g gene_id) using the ‘Ensembl GRCh38.115’ GTF annotation. Ensembl gene identifiers were mapped to gene symbols, gene names, and Entrez IDs using ‘AnnotationDbi’ (v3.21.0; *org.Hs.eg.db*). Differential abundance of genes was analyzed using ‘DESeq2’ (v1.48.2) ^40^ and results used for downstream pathway analyses.

Gene set enrichment analysis (GSEA) was used to evaluate changes in host transcriptional pathways using the Kyoto Encyclopedia of Genes and Genomes (KEGG). A ranked gene list was constructed from DESeq2 generated log2 fold changes after shrinkage using approximate posterior estimation for generalized linear models (*apeglm*) ^41^. Ensembl gene identifiers were mapped to Entrez IDs, and genes without Entrez mapping were excluded. When multiple Ensembl IDs mapped to the same Entrez ID, a single representative entry was retained by selecting the gene with the largest absolute log2 fold change to avoid duplicate identifiers in the ranked list. The signed log2 fold changes were sorted in decreasing order and used as input with analyzed with ‘clusterProfiler’ (v4.16.0; *gseKEGG*, *organism*=“hsa”, *minGSSize*=15, *maxGSSize*=500) ^42^.

### Statistical Analyses

All analyses were performed in R-Studio (*R* v4.4.1). False discovery rate (FDR) was controlled using the Benjamini-Hochberg (BH) procedure for all analyses. For diversity, generalized linear mixed model (GLMM) fit and assumptions were evaluated with ‘DHARMa’ (v0.4.6; *simulateResiduals*, *seed*=8, *n*=250; *testDispersion*; *testOutliers*), and inspection of residuals versus fitted values. For linear mixed models (LMMs), additional diagnostics were conducted with ‘performance’ (v0.15.2; *check_collinearity*; *check_mode; check_normalityl*). Fixed effects were assessed with ‘car’ (v3.1-3; *Anova*, Type III Wald χ^2^). For the primary contrast (fiber intervention; POST vs PRE), model estimates are reported with ‘emmeans’ (v2.0.1) as effect size contrasts (β) with standard errors (SE) and 95% confidence intervals (CI) in **Supplemental Tables**.

### Microbiota diversity and differential abundance (16S rRNA gene

GLMM models for alpha diversity metrics, richness and Shannon, included participant ID as the random intercept with ‘glmmTMB’ (v1.1.9). Richness used a negative binomial distribution (nbinom1) and gaussian for Shannon. Fixed effects included timepoint and baseline covariates (age, sex, BMI, and dietary patterns) with AIC-based comparison used as sensitivity analyses with ‘MASS’ (v7.3-65; *stepAIC*). The ‘vegan’ package was used to test dispersion between timepoints (*betadisper*) and significance of the dispersion (*permutest*, *permutations*=5,000, *pairwise*=T). Community composition differences were tested using permutational multivariate analysis of variance (PERMANOVA) with ‘vegan’ (*adonis2*, *permutations*=5,000, *by*=“margin”) including covariates (age, sex, BMI, history of CRC, history of colonic polyps, smoker status, physical activity level, and dietary intake frequences of red meat, fermented foods, fruit, and whole grains). The R^2^ values indicate proportion of compositional variance explained by each predictor. Differential abundance testing was tested with ‘glmmTMB’ nbinom1 models, comparing a null [*ASV ∼ (1|study_id)*] to a full model [*ASV ∼ timepoint + (1|study_id)*] using likelihood ratio tests (LRT), FDR controlled at 0.25. Effect sizes are reported on the model link (log) scale as timepoint-associated log-fold changes in **Supplemental Tables**.

### Targeted Metabolomics

Batch-corrected values were analyzed using LMM by maximum likelihood with ‘lme4’ (v1.1-35.5; *lmer*) which allows all available observations for each metabolite, any missing values are missing-at-random. The model was used to estimate the fiber effect [*metabolite ∼ timepoint + (1|study_id)*] and evaluated using Satterthwaite’s method with ‘lmerTest’ (v3.1-3; *anova*).

### Gene expression and Pathway enrichment

Differential gene expression analysis was tested using a paired design [*∼ study_id + timepoint*] with ‘DESeq2’ (v1.48.0). Differential expression was tested using the default Wald tests. Log2fold changes were shrunk (*type*=“apeglm”) and statistical significance were based on the results, FDR controlled at 0.05 for genes. GSEA pathways, tested with ‘clusterProfiler’ (v4.16.0), were considered significant if pathway met all following criteria: nominal *P*<0.05, FDR ≤0.25, and absolute normalized enrichment score (|NES|)>1 ^43^.

### Microbiota-metabolite associations and predictive modeling

Correlations between changes (Δ, POST-PRE) in mucosal ASVs and ΔSCFAs were tested using Spearman with ‘Hmisc’ (v5.2-5), FDR controlled at 0.25 within serum and tissue analyzed separately. Random forest (RF) regression models were fit with ‘randomForest’ (v4.7-1.2; *ntree*=500; *seed*=8) to predict ΔSCFAs from ΔASVs; predictive performance summarized by out-of-bag variance explained (OOB R^2^) ^44^. For each RF model, performance was assessed by permutation testing. The response was randomly permuted with 1000 iterations, while the predictor matrix held fixed, each permuted dataset was refit, and OOB R^2^ was recorded. Permutation p-values were computed as *P=(b+1)/(nperm+1)*, where *b* is the number of permuted OOB R^2^ values ≥ the observed ^45^. Permutation p-values across the per-metabolite models were FDR controlled at 0.25.

## Results

### Participant characteristics

The cohort (N=37; **Table 1**) was predominantly white (92% of participants) and male (65%), with a mean BMI in the obesity range (30.7 ± 6.7 kg/m^2^). Red meat was commonly reported 1-2 times per week (56.8%; daily: 27.0%), and processed meat consumed monthly by 94.6% (daily: 5.4%). Other frequently reported foods included vegetables (daily: 77.8%), fruits and juices (daily: 64.9%), and whole grains (≥weekly: 91.7%; **Table 2**).

**Table 1.** Baseline Demographic and Clinical Measures (n=37). Data are n (%). Percentages are calculated on non-missing responses. SD = Standard deviation.

|  |  |
| --- | --- |
| **Age** |  |
| Mean (SD) | 54.0 (15.0) |
| Median [25%,75%] | 53.0 [42.0,65.0] |
| **Gender** |  |
| Female | 13 (35.1) |
| Male | 24 (64.9) |
| **Race** |  |
| Black/African American | 1 (2.8) |
| Other(specify) | 2 (5.6) |
| White | 33 (91.7) |
| Missing | 1 |
| **Ethnicity** |  |
| Hispanic or Latino | 2 (5.6) |
| Non-Hispanic or Latino | 34 (94.4) |
| Missing | 1 |
| **BMI** |  |
| Mean (SD) | 30.7 (6.7) |
| Median [25%,75%] | 30.4 [26.1,33.7] |
| **Alcohol use** |  |
| No | 13 (35.1) |
| Yes | 24 (64.9) |
| **Current/active smoker** |  |
| No | 34 (91.9) |
| Yes | 3 (8.1) |
| **History of colorectal cancer** |  |
| No | 29 (78.4) |
| Yes | 8 (21.6) |
| **History of inflammatory bowel disease** |  |
| No | 34 (91.9) |
| Yes | 3 (8.1) |
| **History of colonic polyps** |  |
| No | 20 (54.1) |
| Yes | 17 (45.9) |
| **History of previous colonic surgeries** |  |
| No | 34 (94.4) |
| Yes | 2 (5.6) |
| Missing | 1 |
| *n (%)* |  |

**Table 2.** Baseline Dietary Habits. Data are n (%). Percentages are calculate on non-missing responses; vegetables and whole grain are based on n=36 (n=10. Intake was self-reported by questionnaire using the following question and serving definition: Describe your red meat consumption (beef, pork, lamb, or veal). (1 portion = 4 ounces or approximately the size of a deck of cards. Examples: 4 oz steak, ¼ lb hamburger); Describe your processed meat consumption. (processed meat = bacon, ham, sausage, salami, pepperoni, hot dogs, spam, bologna); Describe your vegetable consumption; (1 serving of vegetables = 1 cup of raw vegetables or ½ cup of cooked vegetables); Describe your fruit consumption. (1 serving fruit = 1 cup raw or canned fruit, small apple, large banana, medium grapefruit, large orange, 1 cup 100% fruit juice); Describe your whole grain consumption (whole wheat bread, oats, brown rice, quinoa, wheat, barley, farro, millet, buckwheat, couscous, etc.) (1 serving of grains = ½ cup cooked oatmeal, 1 slice bread, ½ cup cooked brown rice, ½ cup cooked whole grain pasta); How often do you eat fermented foods? (Examples of fermented foods include yogurt, kefir, sauerkraut, kimchi, kombucha, tempeh, miso, and buttermilk).

|  |  |
| --- | --- |
| **Red meat** |  |
| None | 1 (2.7) |
| More than 1 portion daily | 1 (2.7) |
| 1 portion daily | 9 (24.3) |
| 1–2 portions per week | 21 (56.8) |
| 1–2 portions per month | 5 (13.5) |
| **Processed meat** |  |
| None | 2 (5.4) |
| Daily | 2 (5.4) |
| 2–3 times per week | 12 (32.4) |
| Once weekly | 9 (24.3) |
| Once monthly | 12 (32.4) |
| **Vegetables** |  |
| None | 0 (0.0) |
| More than 3 servings daily | 1 (2.8) |
| 1–3 servings daily | 27 (75.0) |
| 1–3 servings weekly | 8 (22.2) |
| 1–3 servings monthly | 0 (0.0) |
| Missing | 1 |
| **Fruits** |  |
| None | 0 (0.0) |
| More than 3 servings daily | 3 (8.1) |
| 1–3 servings daily | 21 (56.8) |
| 1–3 servings weekly | 9 (24.3) |
| 1–3 servings monthly | 4 (10.8) |
| **Whole grain** |  |
| None | 1 (2.8) |
| More than 3 servings daily | 1 (2.8) |
| 1–3 servings daily | 17 (47.2) |
| 1–3 servings weekly | 15 (41.7) |
| 1–3 servings monthly | 2 (5.6) |
| Missing | 1 |
| **Fermented foods** |  |
| Never | 4 (10.8) |
| Daily | 7 (18.9) |
| 2–3 times per week | 9 (24.3) |
| Once weekly | 9 (24.3) |
| Once monthly | 8 (21.6) |
| *n (%)* |  |

### Mucosal microbial communities shift modestly with fiber supplementation

First, we examined whether 28d of mixed-fiber would produce microbial community-wide remodeling at the mucosal interface. To examine the impact of fiber supplementation on overall community diversity we began by evaluating impacts on richness and entropy (**Figure 1. A-B**). We find that richness (GLMM: β=0.01, *P*=0.93) and entropy (β=0.04, *P*=0.75) were not significantly impacted by the fiber supplementation. Community composition measured by beta diversity (**Figure 1. C-D**) also did not differ significantly following fiber supplementation; NMDS ordination (stress=0.143) showed overlapping distributions between timepoints. PERMANOVA confirmed no significant fiber effect (adonis2: R^2^=0.01, *P*=0.91). In contrast, multiple baseline host and dietary factors significantly influenced composition; fermented foods (R^2^=0.08, *P*=2.00×10^-4^), age (R^2^=0.06, *P*=2.00×10^-4^), and physical activity (R^2^=0.06, *P*=2.00×10^-4^) had the strongest associations. Additional significant predictors included fruit/juice frequency (R^2^=0.04, *P*=2.00×10^-4^), red meat frequency (R^2^=0.04, *P*=4.00×10^-4^), BMI (R^2^=0.04, *P*=4.00×10^-4^), whole grain frequency (R^2^=0.03, *P*=1.20×10^-3^), history of CRC (R^2^=0.02, *P*=2.00×10^-4^), sex (R^2^=0.02, *P*=4.00×10^-4^), history of colonic polyps (R^2^=0.02, *P*=8.00×10^-4^), and active smoker status (R^2^=0.02, *P*=8.00×10^-3^). The full model explained 42% of the compositional variance.

**Figure 1.**
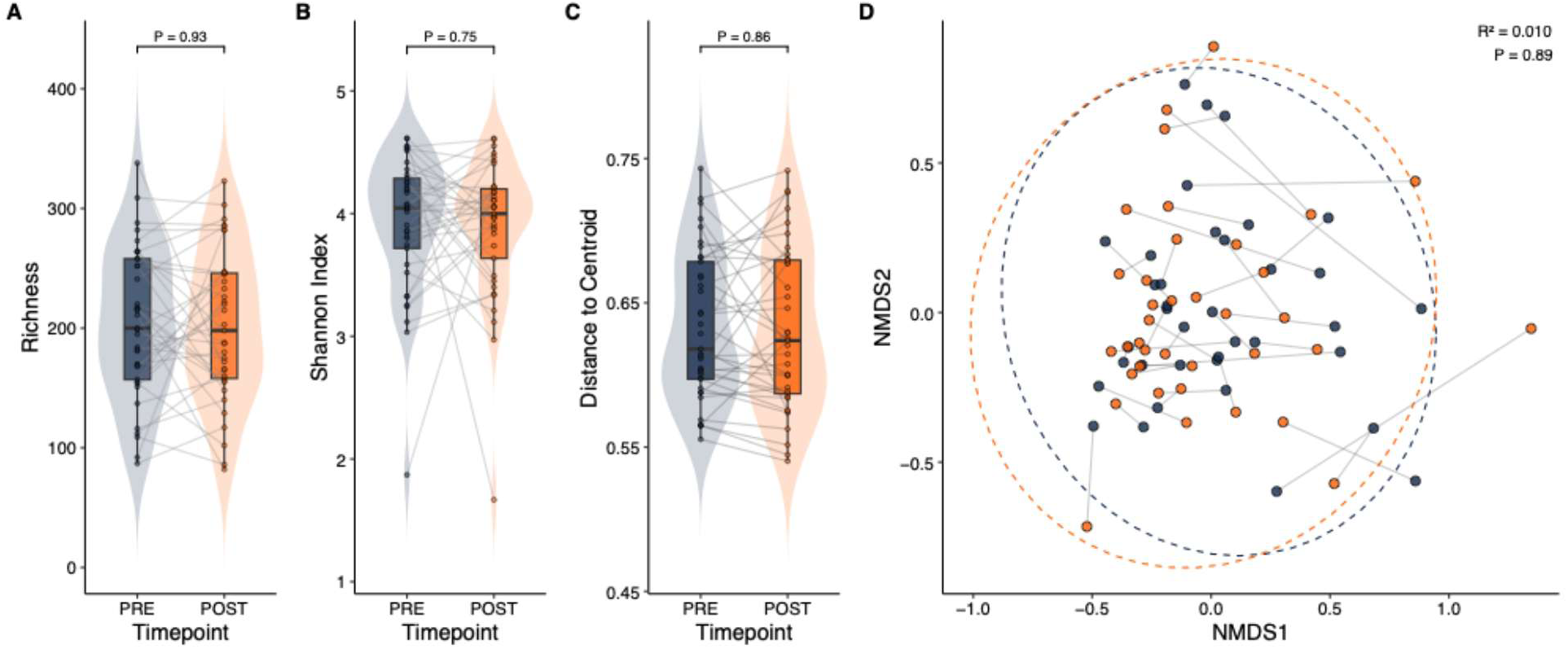
Mixed fiber supplementation effects on 16S bacterial diversity in mucosal tissue. N=37. Alpha diversity (**A**) richness and (**B**) Shannon violin plots show the distribution at both timepoints (PRE=blue, POST=orange), points represent individual samples, and gray lines connect paired measurements within participants. *P*-values are from adjusted generalized linear mixed models (Type III Wald χ^2^ test for timepoint), with participant (ID) modeled as the random intercept. Beta diversity (**C**) distance to centroid based on Jaccard distances (vegan::*betadisper* + permtest) comparing dispersion between timepoints. Beta ordination by (**D**) NMDS using Jaccard distance is colored by timepoint with within-participant pairing indicated by connecting lines; dashed ellipses show 95% confidence regions for each timepoint. Timepoint differences in community composition were tested by PERMANOVA (vegan::*adonis2*, 5,000 permutations; marginal/Type III; adjusted for pre-specified host characteristics and dietary habits); the displayed R^2^ and *P-*value correspond to the timepoint term.

Next, we examined whether the differential abundance of taxa varied with intervention. Fiber supplementation did shift the abundance for 10/302 mucosal ASVs that met the FDR≤0.25 threshold (**Figure 2; Supplemental Table 1**). Notably, we observed increases of ∼6.7-fold and ∼7-fold in two ASVs for *Parabacteroides distasonis* (LRT: ASV89: *P*=1.16×10^-3^, FDR=0.06; ASV152: *P*=1.66×10^-3^, FDR=0.07) and a 2.9-fold and 2.3-fold increase in two ASVs for *Lachnospiraceae* NK4A136 group (ASV144: *P*=4.50×10^-3^, FDR=0.17; ASV130: *P*=5.34×10^-3^, FDR=0.18). Notably, there was a 5.9-fold increase in *Lachnoclostridium* (ASV745; *P*=7.75×10^-4^, FDR=0.06) and 2.4-fold increase in *Bacteroides plebeius* (ASV44; *P*=7.20×10^-3^, FDR=0.22). In contrast, ASVs that decreased were: 1.7-fold in *Coprococcus eutactus* (ASV123: *P*=1.19×10^-4^, FDR=0.04), 0.4-fold in UBA1819 (ASV64: *P*=1.20×10^-3^, FDR=0.06), 3.6-fold in *Blautia* (ASV302: *P*=1.14×10^-3^, FDR=0.06) and 3.3-fold in *Bacteroides eggerthii* (ASV476: *P*=5.78×10^-4^, FDR=0.06). At the genus level, only UBA1819 met the FDR threshold (*P*=1.40×10^-3^, FDR=0.17) with a 0.4-fold decrease (**Supplemental Figure 2**), and none at higher taxonomical ranks (**Supplemental Table 2-5**).

**Figure 2.**
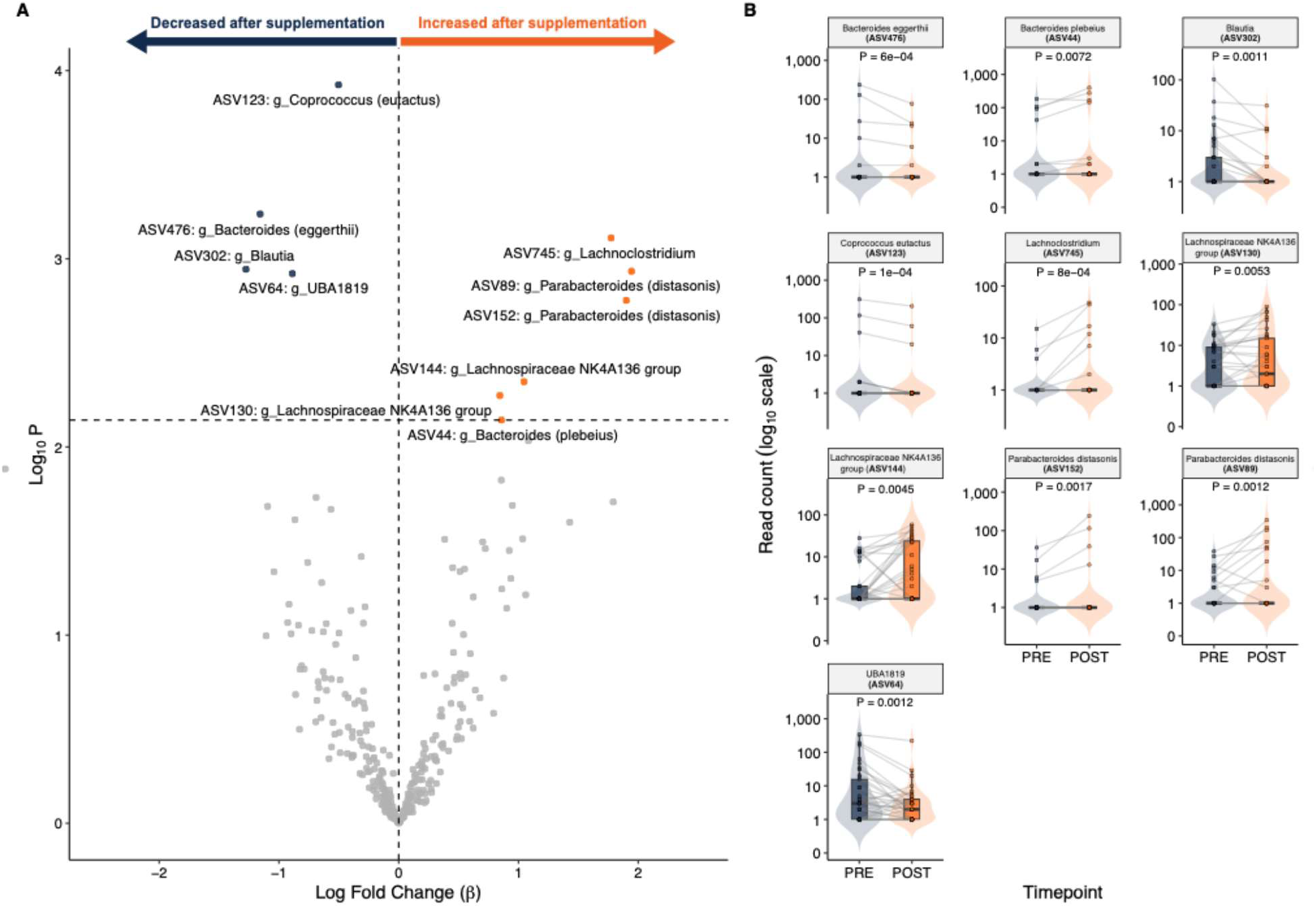
Mixed fiber supplementation effects on 16S differential abundance measured in mucosal tissue. N=37. Differential abundance using ASVs shown as (**A**) volcano plot summarizing generalized linear mixed model fit per ASV (302 ASVs) with participant ID as a random intercept. The x-axis shows the estimated timepoint effect (β; positive values indicate higher relative abundance at POST, negative values indicate lower abundance), and the y-axis shows statistical significance as -log10 of the likelihood ratio test (LRT) *P*-value for the timepoint term. The vertical dashed line marks β = 0; the horizontal dashed line marks the false discovery rate threshold (FDR=0.25). Points are colored by significance and direction (orange, significant increase; blue, significant decrease; gray, not significant). And statistical inferences in (**B**) forest plot show the timepoint effect estimate (POST vs PRE) for ASVs meeting the significance threshold. Points represent the estimated marginal mean contrasts from the fitted mixed-effect model for each ASV (β on the model link scale, interpreted as log-fold change), and horizontal error bars indicate 95% confidence intervals. The dashed vertical line marks no change (β = 0). ASVs are labeled by taxonomy assigned using the SILVA 138.1 classifier with ASV ID in parentheses and ordered by effect size.

### Fiber supplementation alters mucosal short chain fatty acid levels

Since community-wide microbial structure changed only modestly, we next tested whether the mixed-fiber was associated with shifts in downstream fermentation products, quantifying rectal tissue and serum SCFAs as complementary readouts of mucosal and systemic exposure. The metabolome profiles showed a decrease in both tissue and serum (**Supplemental Table 6**). In tissue (**Figure 3. A-E**), propionate resulted with a 1.7-fold decrease (LMM: β=-0.55, *P*=0.04), 2.0-fold decrease in isobutyrate (β=-0.70, *P*=0.03), and a marginal 2.2-fold decrease in acetate (β=-0.79, *P*=0.06). In serum, no changes for either acetate or propionate were observed (all *P*>0.31), however, butyrate had a 1.6-fold decrease (β=-0.49, *P*=3.38×10^-6^) (**Figure 3. F-H**).

**Figure 3.**
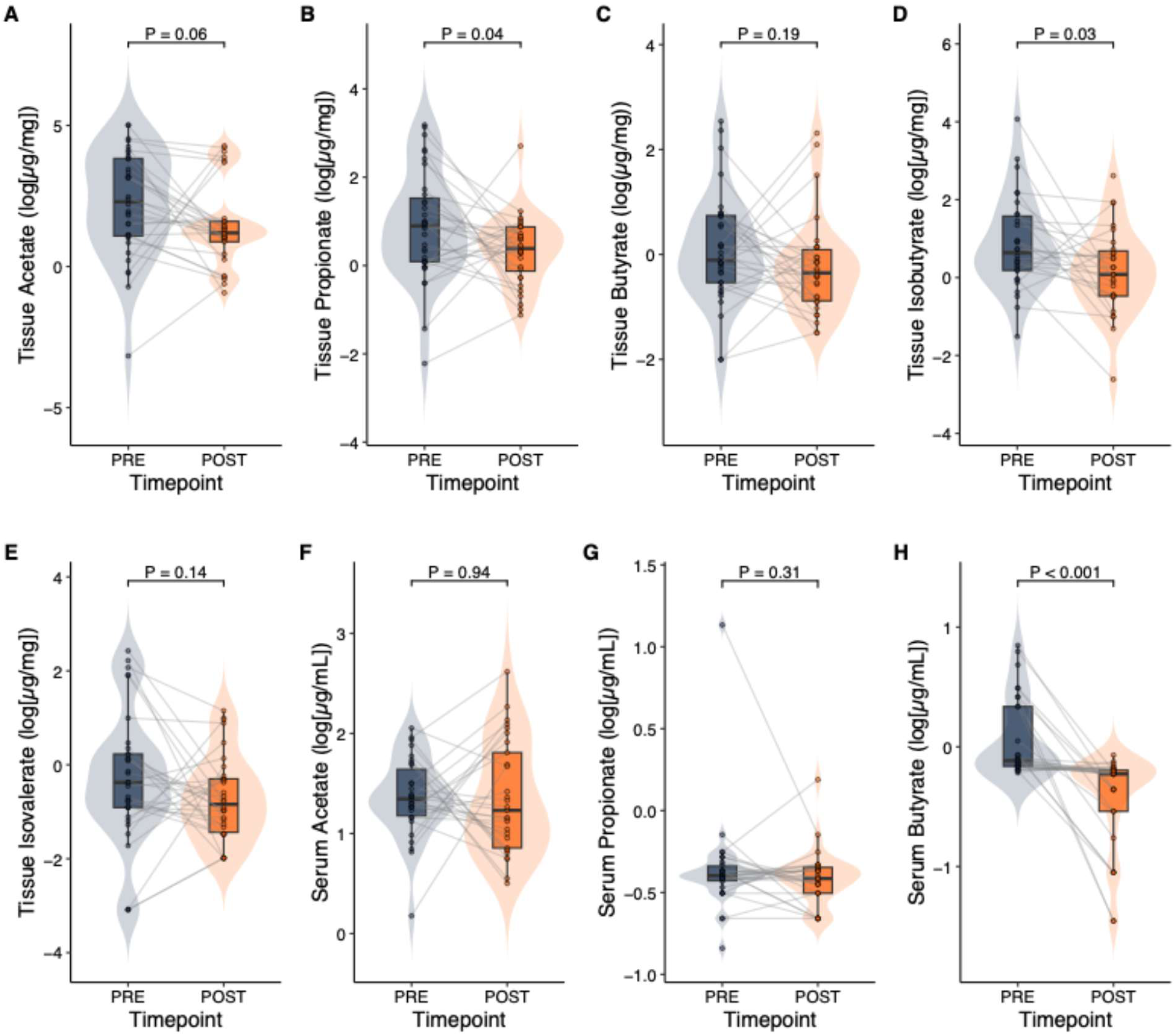
Effects of mixed fiber on targeted metabolomics measured in serum and tissue. N=37. Rectal biopsies were used to measure batch-corrected log-transformed concentrations for short-chain fatty acids (**A-C**) and branch-chain fatty acids (**D-E**) at PRE and POST. And serum SCFAs (**F-H**) were also batch corrected and log transformed. Points represent individual samples and grey lines connect paired observations within participants. P-values correspond the PRE-POST timepoint effect estimated using linear mixed models fit on the batch-corrected log scale.

### Mucosal gene expression shifts towards metabolism with fiber supplementation

Given the changes observed in the microbiota and SCFAs, we next asked if these shifts coincided with altered host mucosal gene expression (**Supplemental Table 7)**. There were 130 differentially expressed genes (DEGs; increased=40, decreased=90; FDR≤0.05 and log2fc≥|0.5|) (**Supplemental Figure 3**). Among DEGs that increased included: 3-hydroxybutyrate dehydrogenase 1 (*BDH1*; *P*=1.98×10^-4^, FDR=0.03), choline dehydrogenase (*CHDH*; *P*=5.19×10^-6^, FDR=0.01), histone deacetylase 11 (*HDAC11*; *P*=1.29×10^-4^, FDR=0.02). Notably, there was concordant directionality in 4-aminobutyrate aminotransferase (*ABAT*; *P*=2.40×10^-3^, FDR=0.09), acyl-CoA dehydrogenase short chain (*ACAD*; *P*=5.40×10^-3^, FDR=0.13), and acyl-CoA synthetase medium-chain family member 3 (*ACSM3*; *P*=2.10×10^-2^, FDR=0.23). In contrast, decreased DEGs included: fibroblast growth factor (*FGF7*; *P*=4.09×10^-8^, FDR=1.71×10^-4^), insulin-like growth factor (*IGF1*; *P*=2.01×10^-6^, FDR=3.35×10^-3^), matrix metallopeptidase 2 (*MMP2*; *P*=5.10×10^-5^, FDR=0.02), toll like receptor 4 (*TLR4*; *P*=3.02×10^-4^, FDR=0.04), interleukin 1 receptor type 1 (*IL1R1*; *P*=1.75×10^-4^, FDR=0.03), and gap junction protein a1/connexin 43 (*GJA1*; *P*=1.75×10^-10^, FDR=2.92×10^-6^).

To contextualize these gene-level shifts at the systems level, we performed GSEA on the pre-ranked transcriptome (**Supplemental Figure 4)**. There were differential changes in 81/346 KEGG pathways (FDR≤0.25, |NES|≥1; **Supplemental Table 8**). Pathways higher at POST consisted of mitochondrial and metabolic programs such as: carbon metabolism (*P*=3.93×10^-7^, FDR= 9.93×10^-5^), oxidative phosphorylation (*P*=1.46×10^-6^, FDR=1.70×10^-4^), biosynthesis of amino acids (*P*=2.36×10^-5^, FDR=1.02×10^-3^), the citric acid cycle (*P*=2.04×10^-4^, FDR=3.93×10^-3^), and pentose phosphate pathway (*P*=6.13×10^-5^, FDR=1.64×10^-3^) (**Figure 4**). SCFA-handling pathways, propanoate metabolism (*P*=4.57×10^-3^, FDR=0.04) and butanoate metabolism (*P*=9.40×10^-3^, FDR=0.06), were also higher at POST (**Supplemental Figure 5-8**). Pathways higher at PRE themed around extracellular matrix/adhesion and innate immune programs including: integrin signaling (*P*=5.73×10^-7^, FDR=9.90×10^-5^), ECM-receptor interaction (*P*=5.99×10^-6^, FDR=3.50×10^-4^), cell adhesion molecule (CAM) interaction (*P*=1.63×10^-5^, FDR=8.10×10^-4^), and complement and coagulation cascades (*P*=6.82×10^-4^, FDR=9.87×10^-3^). Several KEGG disease pathways (e.g., COVID-19 and malaria) were driven by shared inflammatory and complement/coagulation genes rather than a pathogen-specific signature (e.g., *TLR4*, *CGAS*, *CCL2*, *IL6ST*, *C1R*/*C1S*/*C3*/*C5*/*C7*, *CFD*, *MASP1*; *F13A1*). Cell-to-cell communication pathways including cadherin signaling (*P*=1.88×10^-4^, FDR=3.84×10^-3^) and gap junction (*P*=6.10×10^-3^, FDR=4.73×10^-2^), were also higher at PRE with *GJA1* as a dominant leading-edge driver.

**Figure 4.**
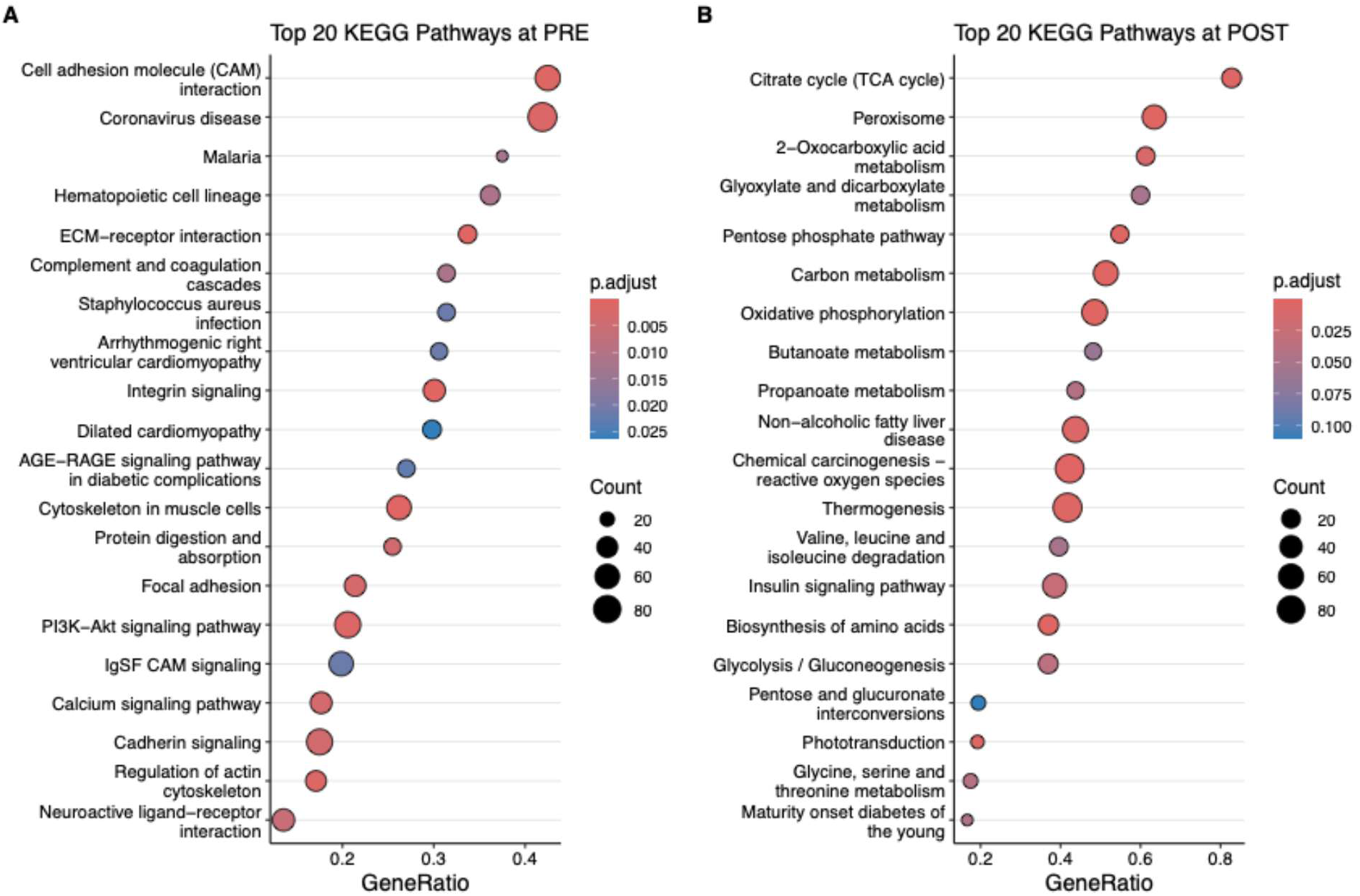
Bidirectional KEGG pathway enrichment in host transcriptome following the fiber intervention. N=10. Preranked KEGG gene set enrichment analysis (GSEA) of host mucosal tissue using RNA-Seq measured pathways enriched in PRE (left; negative NES) and POST (right; positive NES). Panels show the top 20 pathways per direction meeting prespecified significance criteria (P<0.05, FDR<0.25, |NES|>1), ranked by FDR. Gene ratio reflected by point size indicates the number of genes contributing to the pathway signal, and point color the corrected P-value

### Altered mucosal microbial abundance modestly associates with changes in SCFAs

To assess whether intervention-associated shifts in the mucosal microbiota tracked with local SCFA changes, we computed within-participant Δ (POST-PRE) to test spearman correlations between intervention-associated mucosal ΔASVs with ΔSCFAs (**Supplemental Table 9**), and with ΔDEGs (**Supplemental Table 10**). There were no associations between ΔASVs with ΔSCFAs from tissue (all FDR *P*≥0.65; **Supplemental Figure 9a**), ΔASVs with ΔSCFAs from serum (all FDR *P*≥0.84; **Supplemental Figure 9b**), or ΔASVs with ΔDEGs (all FDR≥0.94; **Supplemental Figure 10**).

To widen the microbial scope, mucosal microbiota profiles were used in RF regression models to test whether ΔASV abundances (all 302 ASVs) could predict ΔSCFAs responses (**Supplemental Table 11**). Predictive performance was poor for tissue isobutyrate and propionate (RF: all OOB R^2^<0; **Supplemental Figure 11a, b**) with no evidence of performance beyond a response-permutation null (permutation: *P*=0.98, FDR=0.98; *P*=0.92, FDR=0.98, respectively). In contrast, Δ serum butyrate showed a modest signal (OOB R^2^=0.14) that exceeded the permuted null distribution (*P*=0.02, FDR=4.80×10^-2^), indicating that mucosal community profiles capture a small but non-random component of variance in systemic Δbutyrate. Within the serum butyrate model, the strongest predictors ranked by the percent increase in predictor error (%IncMSE) were ASV60 (*Alistipes*; 7.92%) and ASV51 (*Varibaculum*; 7.26%). Intervention-associated taxons, ASV152 (*Parabacteroides distasonis*; 1.36%) and ASV173 (*Lachnospiraceae* NK4A136 group; 1.43%), were among the top 25 (**Supplemental Figure 11c)**.

## Discussion

In this 28-day mixed-fiber supplementation intervention modeled on how over-the-counter fiber is used in anorectal care, we observed selective shifts in the rectal microbiota, metabolome, and host transcriptome without broad restructuring of community diversity. Community-level alpha and beta diversity remained stable during the intervention, whereas a select subset of mucosal ASVs shifted with supplementation, including increases in ASVs for *Parabacteroides distasonis*, the *Lachnospiraceae* NK4A136 group, and *Lachnoclostridium*. In parallel, and counter to the expectation that increased fiber and SCFA-producing taxa would raise SCFA levels, propionate and isobutyrate in tissue and serum butyrate decreased. The mucosal transcriptome suggested a coordinated functional response; post-intervention KEGG GSEA showed enrichment of intermediary metabolism and SCFA-related pathways, alongside relative attenuation of ECM/adhesion and immune-signaling programs prominent at baseline. Together, these data indicate that short-term fiber supplementation can elicit a coordinated functional response at the mucosal interface, where host-microbe crosstalk is most direct, without broad shifts in community composition, suggesting that functional readouts may be more sensitive to dietary modulation than taxonomy alone.

The rectal mucosa is an ecological niche shaped by proximity to the epithelium, oxygen tension, and host-derived substrates, and provides a more direct window into host-microbiome interactions compared to fecal samples, which reflect the distal luminal environment. Consistent with prior work ^17^, mucosal communities are compositionally distinct from fecal profiles, and mucosal-associated *Lachnospiracea* and *Bacteroides* taxa can carry signals relevant to adenoma status. The ASV-level increases observed here, including *Parabacteroides distasonis* and the *Lachnospiracea* NK4A136 group, fall among health-associated taxa ^46^ and are consistent with a selective, niche-adapted response to the mixed-fiber supplementation rather than a whole community restructuring of the mucosal microbiota. *Lachnospiraceae* are commonly reported as markers of a healthy gut that is depleted during colorectal cancer (CRC) progression ^47,48^. Although nearly half of participants had a personal history of colonic polyps, the design and duration of this study do not permit inference about neoplastic risk; this observation is best read as motivation for future prevention-focused work rather than as evidence of a protective effect. *Lachnospiraceae* are also major SCFA producers, particularly of butyrate ^48^, interestingly here because this taxonomic increase coexisted with reduced measured butyrate. The proposed anti-tumor effect of butyrate by acting (histone deacetylase inhibition, cell-cycle arrest, CD8+ T-cell activation) ^49,50^ lie several inferential steps from the present data and are noted only as context. The genus *Parabacteroides* harbors CAZymes (e.g., GH13, GH43, GH144) capable of degrading resistant dextrin ^51,52^; a clinical trial showed that adults adding daily soluble corn fiber exhibited increased *Parabacteroides*, and parallel *in vitro* work confirmed that *P. distasonis* can ferment this fiber in isolation, without cross-feeding ^53^. This is relevant to our cohort with obesity as *P. distasonis* has been reported to be depleted in populations with obesity ^54^, and its abundance has been associated with reduced obesity-induced systemic inflammation through elevation of metabolites such as nicotinic acid and secondary bile acids, supporting tight-junction integrity and limiting the intestinal permeability that drives mucosal inflammation ^55^. The observed increase in *P. distasonis* ASVs in a cohort with obesity is therefore a favorable cohort-relevant shift. In our cohort, a mucosal *Bacteroides* ASV decreased, consistent with reports that soluble fiber raises *P. distasonis*

while inversely associating with *Bacteroides* abundance ^53^, two closely related genera with overlapping metabolic functions, suggesting niche competition ^51,56^. This direction is notable given that *Bacteroides*, and *B. fragilis* in particular, has been associated with adenomas and CRC ^57–64^. This pattern is consistent with functional redundancy, multiple taxa support overlapping metabolic functions, such that dietary fiber produces changes metabolomic and transcriptional profiles without requiring large shifts in microbial diversity, as observed here.

The second key observation concerns the relationship between SCFA pools and host transcriptional programs after mixed-fiber supplementation. The two fibers ferment differently; psyllium husk is a gel-forming, viscous fiber with slower fermentability that alters luminal hydration, transit, and substrate delivery ^21,22^, whereas wheat dextrin is a soluble, readily fermentable whose α-(1,6/1,4/1,3/1,2) glycosidic bonds are accessible to microbial degradation ^21,65^. This difference shapes where, how quickly, and to what extent SCFAs are generated along the colon. In our cohort, supplementation was associated with decreased tissue propionate and isobutyrate, and decreased serum butyrate, while the mucosal transcriptome showed enrichment of oxidative phosphorylation, intermediary metabolism, and SCFA-related (propanoate and butanoate) pathways. At first past these directions appear discordant, but measured SCFA concentrations reflect a net balance of microbial production, epithelial uptake, and downstream metabolism rather than total output ^66–68^. Low measured SCFA is thus consistent with two non-exclusive explanations, reduced microbial production, or increased epithelial uptake and oxidation, which our data cannot fully distinguish. The concurrent transcriptional enrichment of SCFA-oxidation pathways favors a contribution from increased mucosal utilization rather than reduced production alone; a fraction of locally produced butyrate may be taken up and oxidized by the colonocytes with limited systemic spillover. Within the butanoate pathway, leading-edge genes were enriched for fatty-acid catabolism (e.g., *ABAT*) and β-oxidation genes (e.g., *BDH1*, *ACMS3*, *ACADS*) consistent with increased oxidative-metabolism capacity in the mucosa rather than increased SCFA production. Consistent with a distributed rather than taxon-dominated coupling, RF modeling identified only a modest signal for serum butyrate from mucosal community ASVs, indicating that microbiota-SCFA association here is spread across the community rather than by few responsive-taxa. No physiological endpoint was collected, so these inferences remain transcriptional. Overall, the signatures are consistent with increased colonic metabolic capacity and relative attenuation of mucosal-immune signaling programs, co-occuring with subtle shifts between primary fiber degraders (e.g., *Bacteroides* and *Parabacteroides*) and SCFA-producing secondary consumers (e.g., members of the *Lachnospiraceae* family); our data, however, cannot establish a causal chain among these layers.

This study was designed as a pragmatic, real world, add-on intervention rather than a controlled feeding study, which introduces several limitations. Participants maintained their habitual diet, and background variation in dietary patterns, physical activity, medication use, and other exposures likely diluted detectable intervention effects. The cohort’s clinical heterogeneity – including prior antibiotic exposures, comorbidities, and lifestyle differences – improves translational relevance to routine anorectal care but increases inter-individual variability, which may explain why community-wide diversity remained stable and feature-level signals were modest. Baseline diet was captured by food-frequency reporting, which indicated regular intake of both animal- and plant-derived foods but cannot be translated cleanly into Dietary Guidelines for Americans (DGA) equivalents since cup- and ounce-equivalents vary by food form and density ^69–71^; because baseline diet shapes the fermentable substrate available to the mucosal community, this uncertainty in habitual intake is one plausible contributor to the inter-individual variability we observed in the microbiota and metabolite response. The RNA-seq subset (n=10) improves biological resolution but is underpowered for broad inference; GSEA results should be interpreted as coordinated transcriptional signatures rather than definitive pathway activation. Finally, 16S profiling captures community structure but not strain-level function, CAZyme content, or SCFA-producing capacity; shotgun metagenomics would strengthen claims about functional redundancy and substrate utilization. We acknowledge an alternative reading, that stable diversity, a limited set of differential ASVs, and reduced SCFA pools together indicate minimal effect, however, two features that argue against this is the paired within-subject design controls for the high inter-individual variability of mucosal communities, increasing sensitivity to consistent change. And second, the shifts in taxonomy, metabolome, and transcriptome are mutually coherent rather than isolated, and convergence across independent data layers is harder to attribute to chance than any single layer alone.

## Conclusion

Taken together, within these constraints and despite these limitations, short-term mixed fiber supplementation was associated with selective functional and taxonomic changes at the rectal mucosal interface, where host metabolic and signaling programs directly interface with microbial activity. The dissociation between reduced SCFA pools and enriched SCFA-metabolism pathways described above implies that mucosal functional readouts maybe more sensitive indicators of dietary fiber response than taxonomy alone, particularly in heterogenous clinical cohorts where baseline diet, adiposity, and lifestyles contribute substantial variance. These data do not establish mechanism; rather, they motivate follow-up studies explicitly linking fiber type and dose to microbial carbohydrate utilization, SCFA flux, and epithelial metabolic phenotype and underscore the value of mucosal sampling over stool alone in moving from associations to mechanistic models.

## Acknowledgments

This publication is based on research supported by the Colorectal Cancer Alliance (award: #1013405) and a Cancer Center at Illinois pilot award (award # 9590), and UIUC start-up funds to CAG. This work made use of the equipment, software, and facilities provided by the University of Illinois Urbana Champaign College of Veterinary Medicine Shared Equipment Program’s Biocomputing Shared Resource (BioShaRe). The College of Veterinary Medicine BioShaRe is housed in the Illinois Campus Cluster, a computing resource that is operated by the Illinois Campus Cluster Program (ICCP) in conjunction with the National Center for Supercomputing Applications (NCSA) and which is supported by funds from the University of Illinois at Urbana-Champaign. The Toxicology Scholar award from the University of Illinois at Urbana-Champaign also supported partial funding for this study. And the DNA Services Core Facility at the Roy J. Carver Biotechnology Center (CBC) at University of Illinois Urbana-Champaign for their sequencing services for both DNA and RNA samples.

## Data Availability

Raw sequence data are available in the NCBI Sequence Read Archive (SRA) under the BioProject accession code PRJNA1480771.

## Declaration of generative AI and AI-assisted technologies in the writing process

During the preparation of this manuscript, the author(s) used Grammarly and ChatGPT (OpenAI) for editorial support to improve structure, grammar, and readability of author-written text. After using these tools, the author(s) reviewed and edited as needed and take full responsibility for the content of publication.

## References

1. Foxx-Orenstein AE, Umar SB, Crowell MD. Common Anorectal Disorders. 2014.

2. Gardner IH, Siddharthan R V., Tsikitis VL. Benign anorectal disease: Hemorrhoids, fissures, and fistulas. Ann. Gastroenterol.2020; 33:9–18.

3. Bharucha AE, Knowles CH, Malcolm A. An Evidence-Based Practical Review on Common Benign Anorectal Disorders: Hemorrhoids, Anal Fissure, Dyssynergic Defecation, and Fecal Incontinence. Gastroenterology 2026; 170:50–69.

4. Wald A, Bharucha AE, Limketkai B, Malcolm A, Remes-Troche JM, Whitehead WE, Zutshi M. ACG Clinical Guidelines: Management of Benign Anorectal Disorders. American Journal of Gastroenterology 2021; 116:1987–2008.

5. Nelson RL, Abcarian H, Davis FG, Persky V. 38 Diseases of the Prevalence of Benign Anorectal Disease in a Randomly Selected Population. 1995.

6. Cohee MW, Hurff A, Gazewood JD. Benign Anorectal Conditions: Evaluation and Management. Am Fam Physician 2020; 101;1.

7. Perveen S. Prevalence of Benign Anorectal Diseases: A huge burden on society. Isra Medical Journal [Internet] 2022; 14:12–6. Available from: http://www.imj.com.pk/wp-content/uploads/2022/05/OA-1288-08-21.pdf

8. Alonso-Coello P, Mills E, Heels-Ansdell D, López-Yarto M, Zhou Q, Johanson JF, Guyatt G. Fiber for the Treatment of Hemorrhoids Complications: A Systematic Review and Meta-Analysis CME. Am J Gastroenterol [Internet] 2006; 101:181–8. Available from: www.acg.gi.org/journalcme.

9. Food and Drug Administration (FDA). Food Labeling: Revision of the Nutrition and Supplement Facts Labels: Guidance for Industry - Small Entity Compliance Guide [Internet]. 2020. Available from: http://www.fda.gov/FoodGuidance

10. Food and Drug Administration (FDA). Review of the Scientific Evidence on the Physiological Effects of Certain Non-Digestible Carbohydrates. 2018.

11. Hawkins AT, Davis BR, Bhama AR, Fang SH, Dawes AJ, Feingold DL, Lightner AL, Paquette IM. The American Society of Colon and Rectal Surgeons Clinical Practice Guidelines for the Management of Hemorrhoids. Dis Colon Rectum 2024; 67:614–23.

12. Davids JS, Hawkins AT, Bhama AR, Feinberg AE, Grieco MJ, Lightner AL, Feingold DL, Paquette IM. The American Society of Colon and Rectal Surgeons Clinical Practice Guidelines for the Management of Anal Fissures. Dis Colon Rectum 2023; 66:190–9.

13. Byrd DA, Gomez M, Hogue S, Wan Y, Ortega-Villa A, Warner A, Dagnall C, Jones K, Hicks B, Albert P, et al. Effects of a high-fiber, high-fruit and high-vegetable, low-fat dietary intervention on the rectal tissue microbiome. J Natl Cancer Inst 2025; 117:1237– 44.

14. Makki K, Deehan EC, Walter J, Bäckhed F. The Impact of Dietary Fiber on Gut Microbiota in Host Health and Disease. Cell Host Microbe2018; 23:705–15.

15. Desai MS, Seekatz AM, Koropatkin NM, Kamada N, Hickey CA, Wolter M, Pudlo NA, Kitamoto S, Terrapon N, Muller A, et al. A Dietary Fiber-Deprived Gut Microbiota Degrades the Colonic Mucus Barrier and Enhances Pathogen Susceptibility. Cell 2016; 167:1339–1353.e21.

16. Stene C, Xu J, Fallone de Andrade S, Palmquist I, Molin G, Ahrné S, Thorlacius H, Johnson LB, Jeppsson B. Synbiotics protected radiation-induced tissue damage in rectal cancer patients: A controlled trial. Clinical Nutrition 2025; 49:33–41.

17. Watson KM, Gardner IH, Anand S, Siemens KN, Sharpton TJ, Kasschau KD, Dewey EN, Martindale R, Gaulke CA, Tsikitis VL. Colonic Microbial Abundances Predict Adenoma Formers. Ann Surg 2023; 277:E817–24.

18. Mylonaki M, Rayment NB, Rampton DS, Hudspith BN, Brostoff J. Molecular Characterization of Rectal Mucosa-associated Bacterial Flora in Inflammatory Bowel Disease [Internet]. 2005. Available from: https://academic.oup.com/ibdjournal/article/11/5/481/4683974

19. Lin YF, Sung CM, Ke HM, Kuo CJ, Liu W an, Tsai WS, Lin CY, Cheng HT, Lu MJ, Tsai IJ, et al. The rectal mucosal but not fecal microbiota detects subclinical ulcerative colitis. Gut Microbes 2021; 13:1–10.

20. Mysonhimer AR, Holscher HD. Gastrointestinal Effects and Tolerance of Nondigestible Carbohydrate Consumption. Advances in Nutrition 2022; 13:2237–76.

21. Gibb RD, Sloan KJ, McRorie JW. Psyllium is a natural nonfermented gel-forming fiber that is effective for weight loss: A comprehensive review and meta-analysis. J. Am. Assoc. Nurse Pract.2023; 35:468–76.

22. Hobden MR, Martin-Morales A, Guérin-Deremaux L, Wils D, Costabile A, Walton GE, Rowland I, Kennedy OB, Gibson GR. In Vitro Fermentation of NUTRIOSE® FB06, a Wheat Dextrin Soluble Fibre, in a Continuous Culture Human Colonic Model System. PLoS One 2013; 8.

23. Bliss DZ, Weimer PJ, Jung HJG, Savik K. In vitro degradation and fermentation of three dietary fiber sources by human colonic bacteria. J Agric Food Chem 2013; 61:4614–21.

24. Louis P, Solvang M, Duncan SH, Walker AW, Mukhopadhya I. Dietary fibre complexity and its influence on functional groups of the human gut microbiota. In: Proceedings of the Nutrition Society. Cambridge University Press; 2021. page 386–97.

25. Fernandez-Julia P, Commane DM, van Sinderen D, Munoz-Munoz J. Cross-feeding interactions between human gut commensals belonging to the Bacteroides and Bifidobacterium genera when grown on dietary glycans. Microbiome Research Reports2022; 1.

26. Liu P, Wang Y, Yang G, Zhang Q, Meng L, Xin Y, Jiang X. The role of short-chain fatty acids in intestinal barrier function, inflammation, oxidative stress, and colonic carcinogenesis. Pharmacol. Res.2021; 165.

27. Mansuy-Aubert V, Ravussin Y. Short chain fatty acids: the messengers from down below. Front Neurosci 2023; 17.

28. Hajjar R, Richard CS, Santos MM. The role of butyrate in surgical and oncological outcomes in colorectal cancer. Am. J. Physiol. Gastrointest. Liver Physiol.2021; 320:G601–8.

29. Rahman S, Trone K, Kelly C, Stroud A, Martindale R. All Fiber is Not Fiber. Curr. Gastroenterol. Rep.2023; 25:1–12.

30. Walters W, Hyde ER, Berg-Lyons D, Ackermann G, Humphrey G, Parada A, Gilbert JA, Jansson JK, Caporaso JG, Fuhrman JA, et al. Improved Bacterial 16S rRNA Gene (V4 and V4-5) and Fungal Internal Transcribed Spacer Marker Gene Primers for Microbial Community Surveys. mSystems 2016; 1.

31. Callahan BJ, McMurdie PJ, Rosen MJ, Han AW, Johnson AJA, Holmes SP. DADA2: High-resolution sample inference from Illumina amplicon data. Nat Methods 2016; 13:581–3.

32. Yilmaz P, Parfrey LW, Yarza P, Gerken J, Pruesse E, Quast C, Schweer T, Peplies J, Ludwig W, Glöckner FO. The SILVA and “all-species Living Tree Project (LTP)” taxonomic frameworks. Nucleic Acids Res 2014; 42.

33. Cao Q, Sun X, Rajesh K, Chalasani N, Gelow K, Katz B, Shah VH, Sanyal AJ, Smirnova E. Effects of Rare Microbiome Taxa Filtering on Statistical Analysis. Front Microbiol 2021; 11.

34. Wei R, Wang J, Su M, Jia E, Chen S, Chen T, Ni Y. Missing Value Imputation Approach for Mass Spectrometry-based Metabolomics Data. Sci Rep 2018; 8.

35. Bijlsma S, Bobeldijk I, Verheij ER, Ramaker R, Kochhar S, Macdonald IA, Van Ommen B, Smilde AK. Large-scale human metabolomics studies: A strategy for data (pre-) processing and validation. Anal Chem 2006; 78:567–74.

36. Ritchie ME, Phipson B, Wu D, Hu Y, Law CW, Shi W, Smyth GK. Limma powers differential expression analyses for RNA-sequencing and microarray studies. Nucleic Acids Res 2015; 43:e47.

37. Dobin A, Davis CA, Schlesinger F, Drenkow J, Zaleski C, Jha S, Batut P, Chaisson M, Gingeras TR. STAR: Ultrafast universal RNA-seq aligner. Bioinformatics 2013; 29:15– 21.

38. Liao Y, Smyth GK, Shi W. FeatureCounts: An efficient general purpose program for assigning sequence reads to genomic features. Bioinformatics 2014; 30:923–30.

39. Love MI, Huber W, Anders S. Moderated estimation of fold change and dispersion for RNA-seq data with DESeq2. Genome Biol 2014; 15.

40. Zhu A, Ibrahim JG, Love MI. Heavy-Tailed prior distributions for sequence count data: Removing the noise and preserving large differences. Bioinformatics 2019; 35:2084–92.

41. Yu G, Wang LG, Han Y, He QY. ClusterProfiler: An R package for comparing biological themes among gene clusters. OMICS 2012; 16:284–7.

42. Feng Z, Li Z, Peng W, Zhang J, Li C, Shi R, Li S. RNA-seq analysis identifies key genes and signaling pathways involved in androgen promotion of sebaceous gland proliferation in Hetian sheep. Sci Rep 2025; 15.

43. Breiman L. Random Forests. 2001.

44. Phipson B, Smyth GK. Permutation P-values should never be zero: Calculating exact P-values when permutations are randomly drawn. Stat Appl Genet Mol Biol 2010; 9.

45. Goel A, Shete O, Goswami S, Samal A, C.B. L, Kedia S, Ahuja V, O’Toole PW, Shanahan F, Ghosh TS. Toward a health-associated core keystone index for the human gut microbiome. Cell Rep 2025; 44.

46. Olovo CV, Huang X, Zheng X, Xu M. Faecal microbial biomarkers in early diagnosis of colorectal cancer. J. Cell. Mol. Med.2021; 25:10783–97.

47. Wang H, Zhu W, Lei J, Liu Z, Cai Y, Wang S, Li A. Gut microbiome differences and disease risk in colorectal cancer relatives and healthy individuals. Front Cell Infect Microbiol 2025; 15.

48. Wong CC, Yu J. Gut microbiota in colorectal cancer development and therapy. Nat. Rev. Clin. Oncol.2023; 20:429–52.

49. Lin Y, Lau HCH, Liu C, Ding X, Sun Y, Rong J, Zhang X, Wang L, Yuan K, Miao Y, et al. Multi-cohort analysis reveals colorectal cancer tumor location-associated fecal microbiota and their clinical impact. Cell Host Microbe 2025; 33:589–601.e3.

50. Qu Z, Liu H, Yang J, Zheng L, Huang J, Wang Z, Xie C, Zuo W, Xia X, Sun L, et al. Selective utilization of medicinal polysaccharides by human gut Bacteroides and Parabacteroides species. Nature Communications 2025; 16.

51. Cui Y, Zhang L, Wang X, Yi Y, Shan Y, Liu B, Zhou Y, Lü X. Roles of intestinal Parabacteroides in human health and diseases. FEMS Microbiol. Lett.2022; 369.

52. Alvarado DA, Holthaus TA, Martell S, Southey NL, Atallah M, Sarma R, Revilla D, Brown M, Mehta T, Khan NA, et al. Effects of Soluble Corn Fiber Consumption on Executive Functions and Gut Microbiota in Middle to Older Age Adults: A Randomized Controlled Crossover Trial. J Nutr [Internet] 2026; 156:101473. Available from: https://linkinghub.elsevier.com/retrieve/pii/S0022316626001227

53. Zhang F, Zhang X, Fu J, Duan Z, Qiu W, Cai Y, Ma W, Zhou H, Chen Y, Zheng J, et al. Sex- and Age-Dependent Associations between Parabacteroides and Obesity: Evidence from Two Population Cohort. Microorganisms 2023; 11.

54. Wang K, Liao M, Zhou N, Bao L, Ma K, Zheng Z, Wang Y, Liu C, Wang W, Wang J, et al. Parabacteroides distasonis Alleviates Obesity and Metabolic Dysfunctions via Production of Succinate and Secondary Bile Acids. Cell Rep 2019; 26:222–235.e5.

55. Ezeji JC, Sarikonda DK, Hopperton A, Erkkila HL, Cohen DE, Martinez SP, Cominelli F, Kuwahara T, Dichosa AEK, Good CE, et al. Parabacteroides distasonis: intriguing aerotolerant gut anaerobe with emerging antimicrobial resistance and pathogenic and probiotic roles in human health. Gut Microbes2021; 13.

56. Grion BAR, Fonseca PLC, Kato RB, García GJY, Vaz ABM, Jiménez BN, Dambolenea AL, Garcia-Etxebarria K, Brenig B, Azevedo V, et al. Identification of taxonomic changes in the fecal bacteriome associated with colorectal polyps and cancer: potential biomarkers for early diagnosis. Front Microbiol 2023; 14.

57. Barot S V., Sangwan N, Nair KG, Schmit SL, Xiang S, Kamath S, Liska D, Khorana AA. Distinct intratumoral microbiome of young-onset and average-onset colorectal cancer. EBioMedicine 2024; 100.

58. Chen F, Dai X, Zhou CC, Li KX, Zhang YJ, Lou XY, Zhu YM, Sun YL, Peng BX, Cui W. Integrated analysis of the faecal metagenome and serum metabolome reveals the role of gut microbiome-associated metabolites in the detection of colorectal cancer and adenoma. Gut 2022; 71:1315–25.

59. Dulal S, Keku TO. Gut microbiome and colorectal adenomas. Cancer Journal (United States)2014; 20:225–31.

60. Kim Y, Kim MK, Lee S. Comparative microbiome analysis of paired mucosal and fecal samples in Korean colorectal cancer patients. Front Oncol 2025; 15.

61. Torshizi Esfahani A, Zafarjafarzadeh N, Vakili F, Bizhanpour A, Mashaollahi A, Karimi Kordestani B, Baratinamin M, Mohammadpour S. Gut microbiome in colorectal cancer: metagenomics from bench to bedside. JNCI Cancer Spectr.2025; 9.

62. Zwezerijnen-Jiwa FH, Sivov H, Paizs P, Zafeiropoulou K, Kinross J. A systematic review of microbiome-derived biomarkers for early colorectal cancer detection. Neoplasia (United States) 2023; 36.

63. Shen F, Xu C, Wang C. Gut Microbiome Diagnostic Biomarkers for Colorectal Cancer. Turkish Journal of Gastroenterology 2025;

64. Slavin JL, Savarino V, Paredes-Diaz A, Fotopoulos G. A Review of the Role of Soluble Fiber in Health with Specific Reference to Wheat Dextrin. 2009.

65. Scientific Opinion on the substantiation of health claims related to “wheat dextrin” and maintenance of normal blood pressure (ID 844, 1682), maintenance of normal (fasting) blood concentrations of triglycerides (ID 844, 1682), maintenance of normal blood. EFSA Journal [Internet] 2010; 8:1761. Available from: http://doi.wiley.com/10.2903/j.efsa.2010.1761

66. U.S. Department of Health and Human Services and U.S. Department of Agriculture. 2015-2020 Dietary Guidelines for Americans [Internet]. 2015. Available from: https://odphp.health.gov/dietaryguidelines/2015/guidelines/.

67. Dietary Guidelines for Americans, 2025–2030: Daily Servings by Calorie Level. 2025.

68. Guidelines Advisory Committee D. Scientific Report of the 2020 Dietary Guidelines Advisory Committee Advisory Report to the Secretary of Agriculture and Secretary of Health and Human Services.

69. Den Besten G, Van Eunen K, Groen AK, Venema K, Reijngoud DJ, Bakker BM. The role of short-chain fatty acids in the interplay between diet, gut microbiota, and host energy metabolism. J. Lipid Res.2013; 54:2325–40.

